# A glycosaminoglycan extract from Portunus pelagicus inhibits BACE1, the β secretase implicated in Alzheimer’s disease

**DOI:** 10.1101/613695

**Authors:** Courtney J. Mycroft-West, Lynsay C. Cooper, Anthony J. Devlin, Patricia Procter, Scott Eric Guimond, Marco Guerrini, David G. Fernig, Marcelo A. Lima, Edwin A. Yates, Mark A. Skidmore

## Abstract

Therapeutic options for Alzheimer’s disease, the most common form of dementia, are currently restricted to palliative treatments. The glycosaminoglycan heparin, widely used as a clinical anticoagulant, has previously been shown to inhibit the Alzheimer’s disease-relevant β-secretase 1 (BACE1). Despite this, the deployment of pharmaceutical heparin for the treatment of Alzheimer’s disease is largely precluded by its potent anticoagulant activity. Furthermore, ongoing concerns regarding the use of mammalian sourced heparins, primarily due to prion diseases and religious beliefs, hinder the deployment of alternative heparin based therapeutics. A marine-derived, heparan sulphate-containing glycosaminoglycan extract isolated from the crab *Portunus pelagicus*, was identified to inhibit human BACE1 with comparable bioactivity to that of mammalian heparin (IC_50_ = 1.85 μg.mL^-1^ (R^2^ = 0.94) and 2.43 μg.mL^-1^ (R^2^ = 0.93), respectively) possessing highly attenuated anticoagulant activities. The results from several structural techniques suggest that the interactions between BACE1 and the extract from *P. pelagicus* are complex and distinct from those of heparin.

## 1. Introduction

Alzheimer’s disease (AD), the most common form of dementia, is characterized by progressive neurodegeneration and cognitive decline [1]. The pathological hallmarks of AD include the accumulation of extracellular β-amyloid plaques and intraneuronal neurofibrillary tangles (NFTs) [2]. Deposition and aggregation of toxic amyloid-β proteins (Aβ), the primary constituents of β-amyloid plaques, has been identified as one of the primary causative factors in the development of AD. Approximately 270 mutations within genes that are directly associated with Aβ production are currently linked to the early-onset development of AD [3].

Amyloid-β peptides (Aβ) are produced through the sequential cleavage of the type 1 transmembrane protein, amyloid precursor protein (APP). APP is initially cleaved by the aspartyl protease, β-site amyloid precursor protein cleaving enzyme 1 (BACE1), the primary neuronal β-secretase [4], liberating a soluble N-terminal fragment (sAPPβ) and a membrane bound C-terminal fragment (β-CTF or C99). The β-CTF/C99 fragment subsequently undergoes cleavage by γ-secretase within the transmembrane domain, releasing a 36-43 amino acid peptide (Aβ) into the extracellular space; the most predominant species of Aβ being Aβ40 [5,6]. An imbalance favouring the production of Aβ42 has been linked to the development of AD owing to a higher propensity to oligomerize and form amyloid fibrils, than the shorter Aβ40 [7].

As the rate-limiting step in Aβ production, inhibition of BACE1 has emerged as a key drug target for the therapeutic intervention of the progression of AD, in order to prevent the accumulation of toxic Aβ [8,9]. This is supported by the finding that BACE1 null transgenic mice models survive into adulthood with limited phenotypic abnormalities while exhibiting a reduction in brain Aβ levels [4,10–15]. Despite the therapeutic potential of BACE1 inhibition, the successful development of a clinically approved pharmaceuticals has proven a challenge due to the large substrate-binding cleft of BACE1, and unfavourable *in vivo* pharmaceutical properties of potent peptide inhibitors, for example oral bioavailability, half-life and blood brain barrier (BBB) penetration [9,16].

Heparan sulphate (HS), and its highly sulfated analogue heparin (Hp), are members of the glycosaminoglycan (GAG) family of linear, anionic polysaccharides. They share a repeating disaccharide backbone consisting of a uronic acid (D-glucuronic acid; GlcA or L-iduronic acid; IdoA) and D-glucosamine, which can be variably sulphated or N-acetylated. HS is synthesised attached to a core protein forming HS proteoglycans (HSPGs), which have been identified colocalized with BACE1 on cell surfaces, in the Golgi complex and in endosomes [17]. HSPGs were reported to endogenously regulate BACE1 activity in *vivo* through either a direct interaction with BACE1 and/or by sequestration of the substrate APP [17]. Addition of exogenous HS or heparin was also shown to inhibit BACE1 activity in *vitro* and reduced the production of Aβ in cell culture (17–19). Mouse models treated with low molecular weight heparin (LMWH) exhibit a reduction in Aβ burden [20] and display improved cognition [21]. Furthermore, heparin oligosaccharides within the minimum size requirement for BACE1 inhibition [17,18], (<18-mers) possess the ability to cross the blood brain barrier (BBB) [22] and can be made orally bioavailable, depending on formulation and encapsulation methods [23]. Heparin analogues, therefore hold therapeutic potential as a treatment against AD, which may also offer an advantage over small molecule and peptide inhibitors of BACE1.

Heparin has been utilized clinically as a pharmaceutical anticoagulant for over a century due to its ability to perturb the coagulation cascade, principally through interactions with antithrombin III via the pentasaccharide sequence [-4-α-D-GlcNS,6S (1-4) β-D-GlcA (1-4) α-D-GlcNS,3S,6S (1-4) α-L-IdoA2S (1-4) α-D-GlcNS,6S. The side-effect of anticoagulation presents as an important consideration when determining the potential of a heparin-based pharmaceutical for the treatment of AD. It has been previously determined that the anticoagulation potential of heparin can be highly attenuated by chemical modifications, while retaining the favourable ability to inhibit BACE1 [17–19]. Polysaccharides in which the 6-O-sulfate had been chemically removed were reported to have attenuated BACE1 activity [17,18] although this correlates with an augmented rate of fibril formation [24].

Polysaccharides analogous to GAGs have been isolated from a number of marine invertebrate species that offer rich structural diversity and display highly attenuated anticoagulant activities compared to mammalian counterparts [25–33]. The largely unexplored chemical diversity of marine derived GAGs, provides a vast reservoir for the discovery of novel bioactive compounds, some of which have been identified to exhibit antiviral [34,35], anti-parasitic [36,37], anti-inflammatory [38,39], anti-metastasis [30-41], anti-diabetic [42], anti-thrombotic [32] and neurite outgrowth-promoting activities [43]. Also, these compounds may be obtained from waste material, which makes their exploitation both economical and environmental appealing. Here, a GAG extract isolated from the crab *Portunus pelagicus*, has been found to possess attenuated anticoagulant activity, whilst potently inhibiting the AD relevant β-secretase, BACE1 *in vitro*.

## 2 Results

### 2.1. Isolation and characterisation of a glycosaminoglycan extract from the crab Portunus pelagicus

A glycosaminoglycan extract isolated from the crab *Portunus pelagicus* via proteolysis was fractionated by DEAE-Sephacel anion-exchange chromatography utilizing a stepwise sodium chloride gradient. The eluent at 1 M NaCl (fraction 5; designated *P. pelagicus* F5) was observed to have similar electrophoretic mobility in 1,3-diaminopropane buffer (pH 9.0) to mammalian HS/Hp, with no bands observed corresponding to monosulphated chondroitin sulphate (CSA/CSC), disulphated chondroitin sulphate (CSD) or dermatan sulphate (DS) (Figure 1).

**Figure 1:**
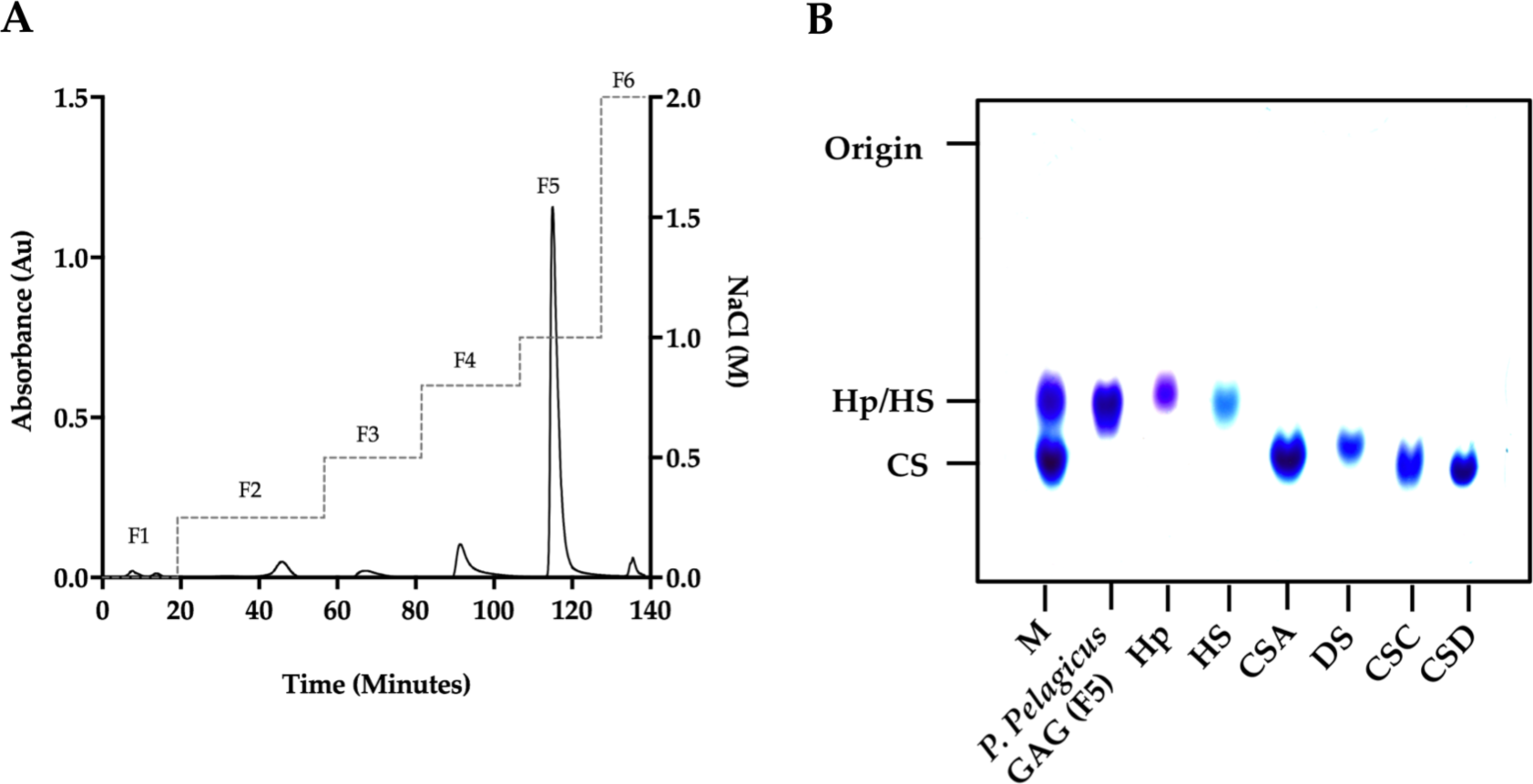
**(A)** DEAE purification of *P. pelagicus* crude glycosaminoglycan. Fractions 1-6 (F1-6; λ_Abs_ = 232 nm, solid line) were eluted using a stepwise NaCl gradient with HPAEC (dashed line). **(B)** Agarose gel electrophoresis of *P. pelagicus* F5. The electrophoretic mobility of *P. pelagicus* F5 was compared to that of *bone fide* glycosaminoglycan standards, heparin (Hp;), heparan sulphate (HS;), dermatan sulphate (DS;) and chondroitin sulphate A, C and D (CSA, CSC and CSD, respectively;). M: CSA, Hp and HS mixture.

In order to corroborate the Hp/HS like structural characteristics of *P. pelagicus* F5, the ATR-FTIR spectra has been compared with that of Hp. Both *P. pelagicus* F5 and Hp were shown to share similar spectral features, for instance bands at 1230, 1430 and 1635 cm^-1^, which are associated with S=O stretches, symmetric carbonyl stretching and asymmetric stretches respectively, indicative of common structural motifs. An additional peak and a peak shoulder located at ∼ 1750 and ∼ 1370 cm^-1^, were observed in *P. pelagicus* F5, but absent in Hp. The peak shoulder at ∼1370 cm^-1^ is indicative of a Hp and CS mixture. The differences observed between the spectra of *P. pelagicus* F5 and Hp in the variable OH region (> 3000 cm^-1^) are likely to be associated with changeable moisture levels present during sample acquisition (Figure 2A) as opposed to underlying differences within glycan structure [44].

**Figure 2:**
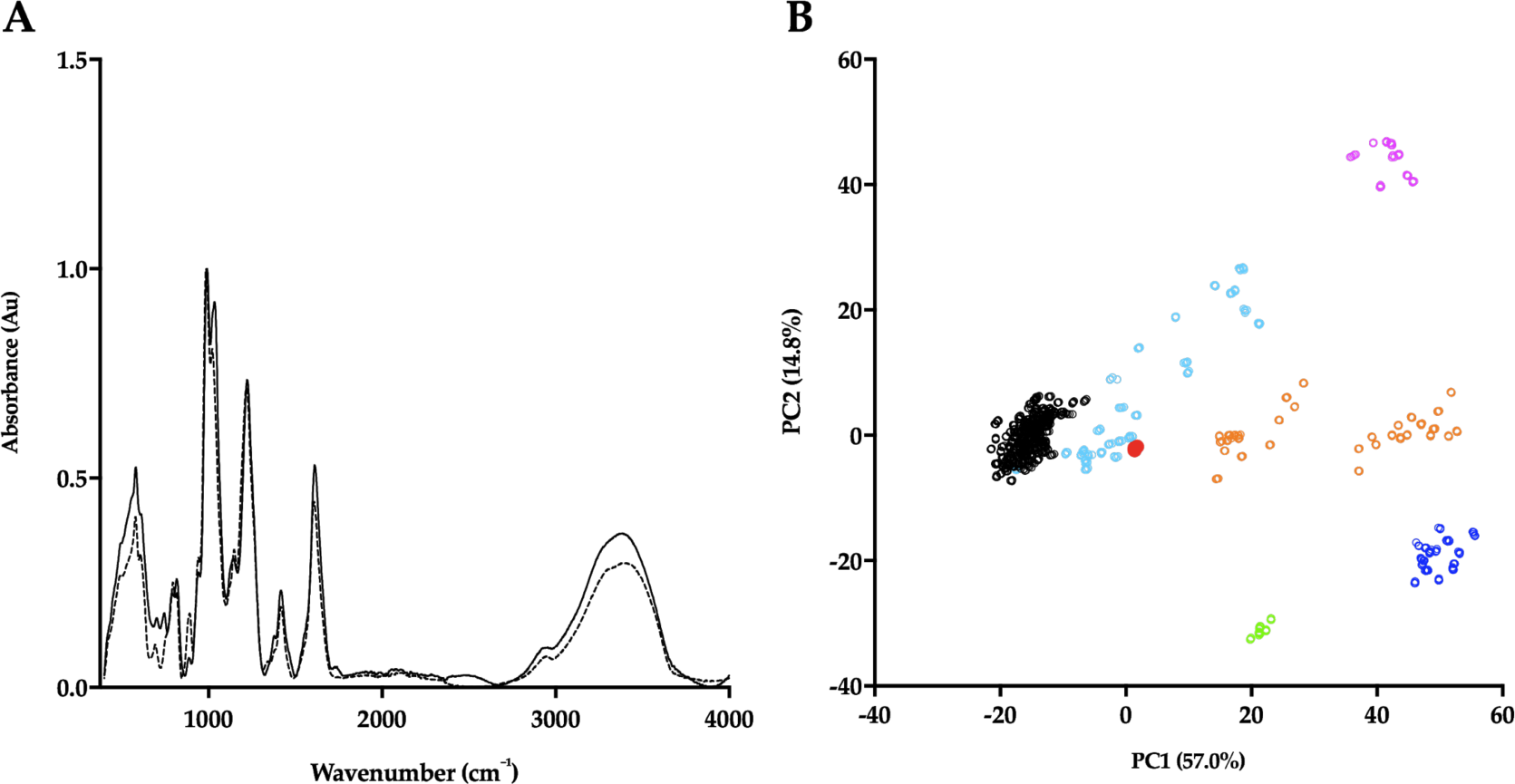
**(A)** ATR-FTIR spectra of porcine mucosal Hp (dashed line) and *P. pelagicus* F5; (solid line), n = 5. **(B)** Principal component analysis (PCA) Score Plot for PC1 vs PC2 of *P. pelagicus* F5 against a *bone fide* GAG library. Hp, black; HS, cyan; CS, orange; DS, blue; hyaluronic acid (HA), magenta; oversulphated-CS, light green and *P. pelagicus* F5, red (filled circle).

Post acquisition, the ATR-FTIR spectrum of *P. pelagicus* F5 was interrogated against a library of known GAGs comprising; 185 Hps, 31 HSs, 44 CSs & DSs, 11 hyaluronic acids (HAs) and 6 oversulfated chondroitin sulphates (OSCSs) using principal component analysis (PCA) [44]. Principal component 1 (PC1), which covers 57% of the total variance, indicates that *P. pelagicus* F5 locates within the region containing mammalian Hp/HS. Through comparison of PC1 and PC2, comprising > 70% of the total variance, *P. pelagicus* F5 lies towards the CS region, a location previously identified with Hps containing small amounts of CS/DS [44], analogous to crude, pharmaceutical Hp.

*P. pelagicus* F5 was subsequently subjected to exhaustive enzymatic cleavage with *Flavobacterium heparinum* lyases I, II and III. The digest products from *P. pelagicus* F5 (Figure 4, Table 1) and a Hp control (Figure 3, Table 1) were analyzed using strong anion-exchange chromatography and the retention times compared to those of the eight common Δ-disaccharide standards present within both Hp and HS [45].

**Table 1:** Corrected disaccharide composition analysis of *P. pelagicus* F5 and Hp.

| Δ-Disaccharide | *P. pelagicus* F5 (%) | Hp (%) |
| --- | --- | --- |
| Δ-UA-GlcNAc | 2.8 | 4.3 |
| Δ-UA-GlcNS | 5.6 | 4.2 |
| Δ-UA-GlcNAc(6S) | 2.3 | 5.0 |
| Δ-UA(2S)-GlcNAc | 16.5 | 3.1 |
| Δ-UA-GlcNS(6S) | 20.2 | 22.9 |
| Δ-UA(2S)-GlcNS | 23.5 | 5.9 |
| Δ-UA(2S)-GlcNAc(6S) | 6.0 | 3.1 |
| Δ-UAs(2S)-GlcNS(6S) | 23.1 | 51.5 |

**Figure 3:**
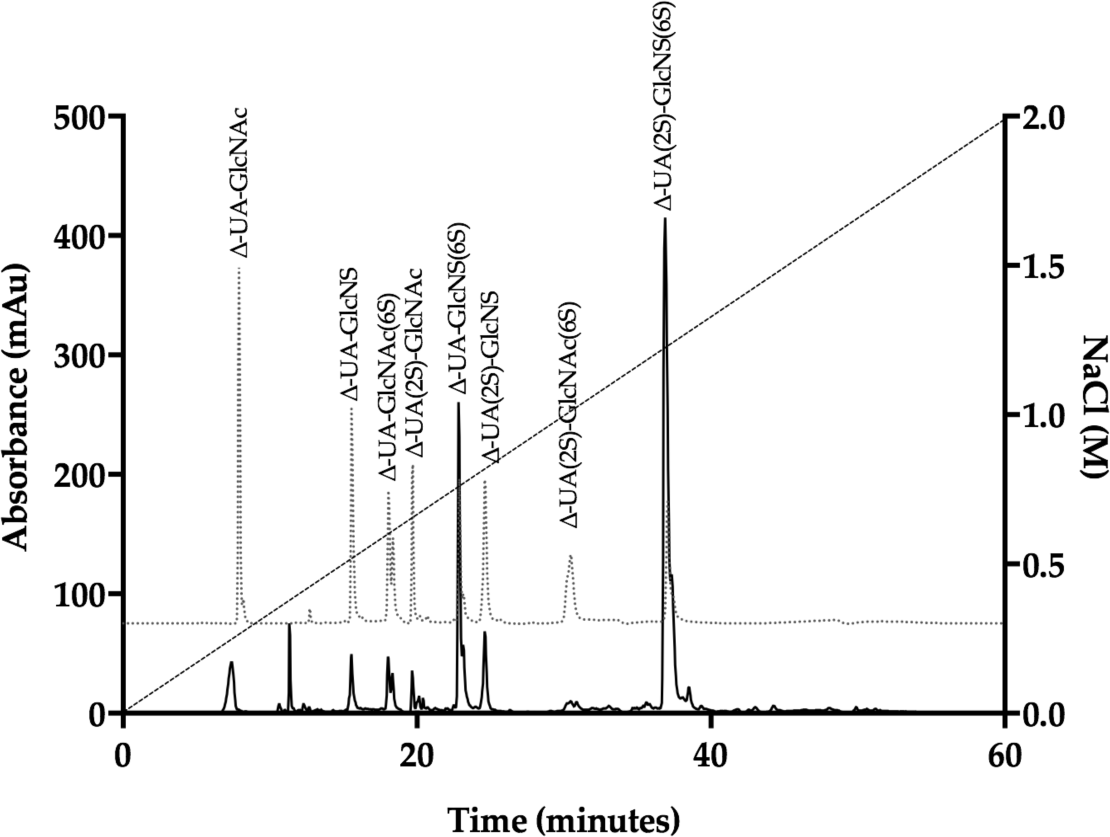
UV-SAX HPLC disaccharide composition analysis was performed on the bacterial lyase digest of Hp (λ_Abs_ = 232 nm) eluting with a linear gradient of 0 - 2 M NaCl (dashed line). Eluted Δ-disaccharides were referenced against the eight common standards present within Hp and HS (light grey, dotted line).

**Figure 4:**
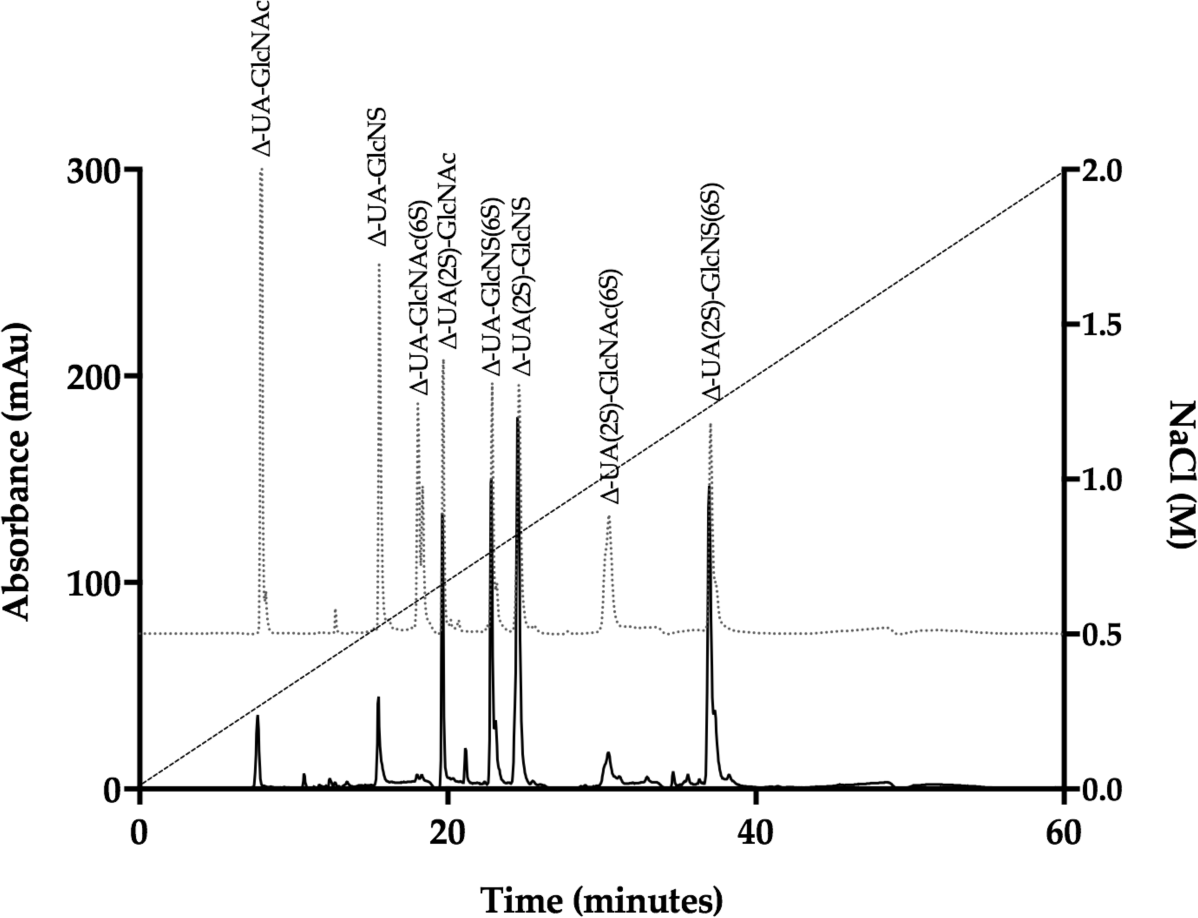
UV-SAX HPLC disaccharide composition analysis was performed on the bacterial lyase digest of the *P.pelagicus* F5 (λ_Abs_ = 232 nm) eluting with a linear gradient of 0 - 2 M NaCl (dashed line). Eluted Δ-disaccharides were referenced against the eight common standards present within Hp and HS (light grey, dotted line).

The digest products detected for Hp were in agreement with a typical mammalian Hp disaccharide profile [45], with 51.5% of the total products attributable to the trisulphated, Δ-UA(2S)-GlcNS(6S) and 22.9% to Δ-UA-GlcNS(6S). A minimal proportion of mono or unsulphated disaccharides, accounting for 12.3%, and 4.3% respectively, were also observed for Hp. In comparison, a more disperse sulphation profile was observed for *P. pelagicus* F5 than Hp (Table 1), with a comparatively lower proportion of trisulphated disaccharides, 23.1%. The *P. pelagicus* F5 contained 24.4% monosulphated disaccharides, of which 16.5% was accounted for by Δ-UA(2S)-GlcNAc. A higher proportion of Δ-UA(2S)-GlcNS (23.5%), was also detected in *P. pelagicus* F5, than Hp (5.9%), indicating that the compound displays distinct structural characteristics. Such features also contrast with that of HS, where ∼50-70% of disaccharides are comprised of Δ-UA-GlcNAc/Δ-UA-GlcNS [25,45–47]. Also, *P. pelagicus* F5 presents a significant higher proportion of trisulphated disaccharides than commonly present in mammalian HS, a typical marker of more heparin-like structures.

Proton and Heteronuclear Single-Quantum Correlation (HSQC) NMR was employed to confirm the GAGs composition of *P. pelagicus* F5. ^1^H NMR can indicate the major signals associated with HS as well as signals that arise from galactosaminoglycans such as CS. The presence of both (Figure 5A insert) is easily identified by the two N-acetyl signals at 2.02 ppm (CS) and 2.04 ppm (HS). ^1^H-^13^C HSQC NMR (Figure 5B) has been used to resolve overlapping signals and saccharide composition estimates using peak volume integration. Integration of N-acetyl signals revealed that the extract is composed of approximately 60% HS and 40% CS. The combined integration of the N-acetyl and A2 signals from the HS showed that *P. pelagicus* F5 possesses a high NS content of approximately 76%, which supports the HPLC-based empirical disaccharide analysis (Figure 4 and Table 1). Together, these data establish that the HS of *P. pelagicus* F5 is considerably more sulfated (Table 1) than that commonly extracted from mammalian sources [45]. With regard to the CS element of *P. pelagicus* F5, signals typical of the CS backbone are present although sulfation is generally low, with galactosamine 6-O-sulfation occurring in approximately 35% of all CS residues. The lack of non-overlapping signal for galactosamine 4-O-sulfation indicates all but negligible levels of such a modification are present within the CS component.

**Figure 5:**
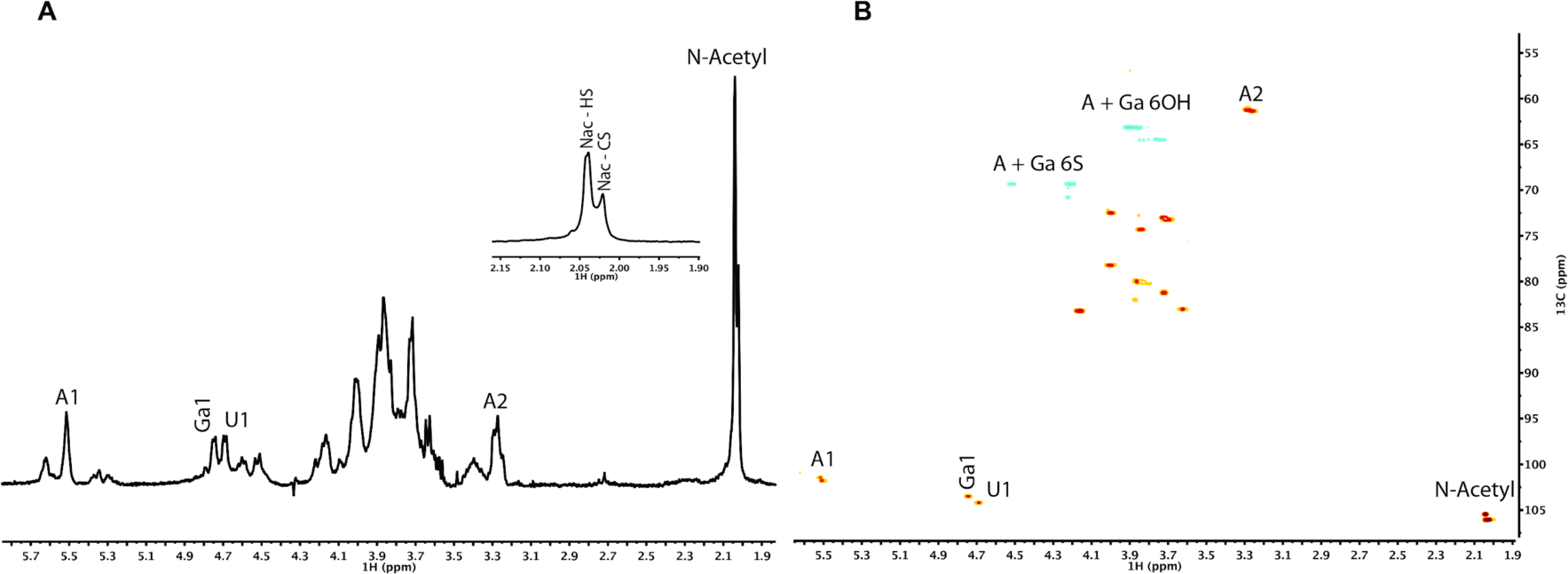
**(A)** ^1^H and **(B)** ^1^H-^13^C HSQC NMR spectra of *P. pelagicus* F5. Major signals associated with HS and CS are indicated. Spectral integration was performed on the HSQC using labeled signals. Key: glucosamine, A; uronic acid, U ; N-Acetyl, Nac and galactosamine, Ga.

### 2.2. P. pelagicus F5 inhibits the Alzheimer’s Disease β-secretase 1

*P. pelagicus* F5 was assayed for inhibitory potential against BACE1, utilizing a fluorogenic peptide cleavage FRET assay, based upon the APP Swedish mutation. Reactions were performed at pH 4.0, within the optimal pH range for BACE1 activity (Figure 6). A maximal level of BACE1 inhibition of 90.7% ± 2.9 (n = 3) was observed in the presence of 5 μg.mL^-1^ *P. pelagicus* F5, with an IC_50_ value of 1.9 μg.mL^-1^ (R^2^ = 0.94). This was comparable to the activity of Hp, which exhibited a maximal level of BACE1 inhibition of 92.5% ± 1.5 (n = 3) at 5 μg.mL^-1^, with an IC_50_ of 2.4 μg.mL^-1^ (R^2^ = 0.93).

**Figure 6:**
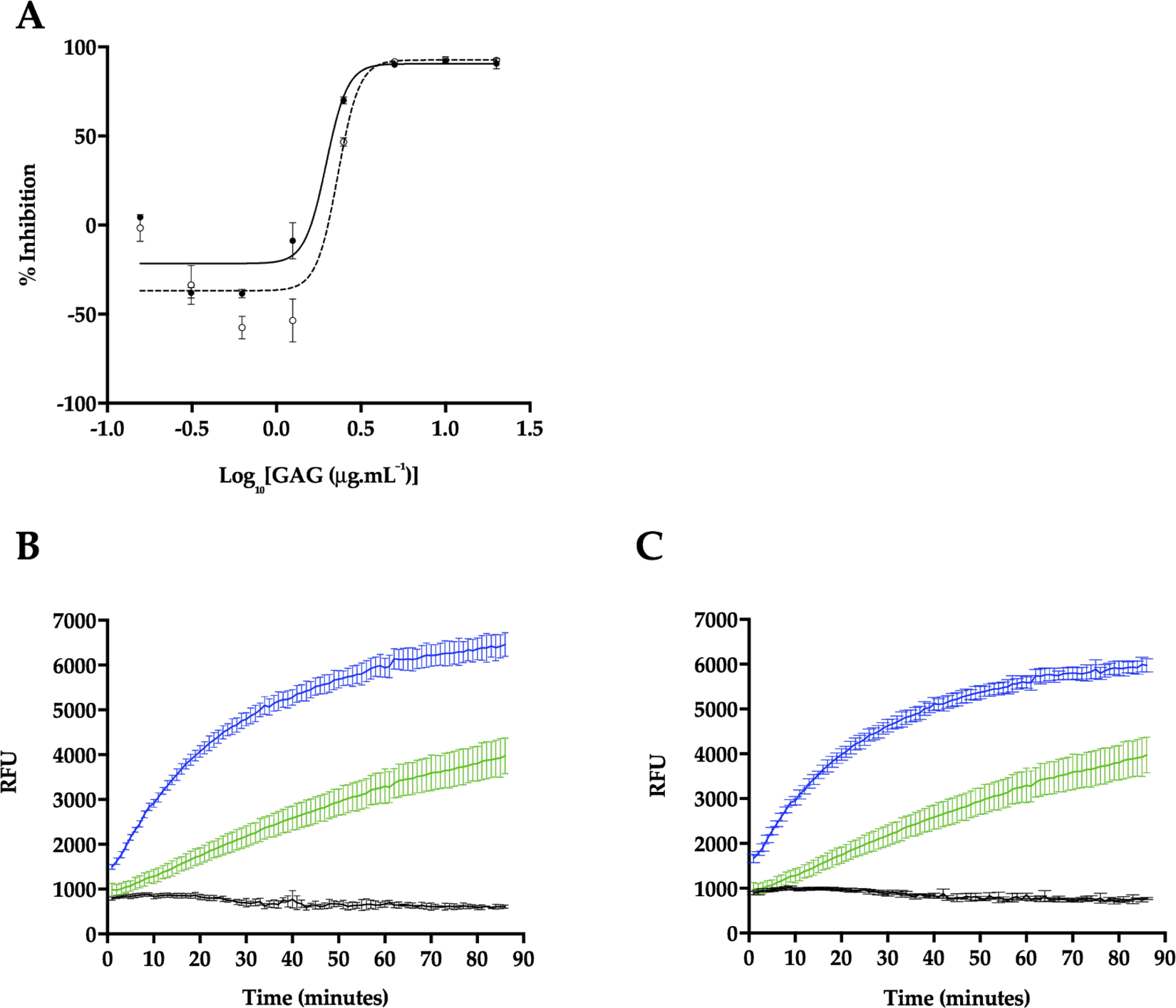
Inhibition of human BACE1 by Hp or *P. pelagicus* F5. **(A)** Dose response of Hp (dashed line, open circles) or *P. pelagicus* F5 (solid line, filled circles) as determined using FRET. *P. pelagicus* F5, IC_50_ = 1.9 μg.mL^-1^ (R^2^ = 0.94); Hp, IC_50_ = 2.4 μg.mL^-1^ (R^2^ = 0.93). **(B)** Time course activation or inhibition of BACE1 by 5 μg.mL^-1^(black) or 625 ng.mL^-1^ (blue)) Hp, compared to water control (green). **(C)** The same as (B) for *P. pelagicus* F5.

In the presence of low concentrations of Hp, an increase in BACE1 activity was observed (Figure 6A-B), with maximal activation occurring at 625 ng.mL^-1^ (57.5% ± 3.7, n = 3). The BACE1 utilised in this study consisted of the zymogen form (Thr^22^-Thr^457^), containing the prodomain sequence. This is in accord with previous reports that demonstrate low concentrations (∼1 μg.mL^-1^) of heparin can stimulate proBACE1 activity [48,49]. A maximum increase in BACE1 activity was also detected in the presence of 625 ng.mL^-1^ of *P. pelagicus* F5 (38.5% ± 1.4, n = 3), although significantly diminished promotion was displayed compared to the same concentration of Hp (57.5% ± 3.7, n = 3); t(4) = 4.859, p = 0.0083. This indicates that although *P. pelagicus* F5 exhibits stimulatory activity, it is significantly less than that of Hp. The percent activity level returned to that of the negative control value at concentrations lower than 312.5 ng.mL^-1^, indicating that both inhibitory and stimulatory effects are dose dependent. For both Hp and *P. pelagicus* F5, BACE1 promotion was followed by enzyme inhibition, as previously reported (Figure 6B-C;[49]. The rate of BACE1 activity between 60 - 90 minutes was significantly different from controls lacking either Hp (n = 3-6; t(4) = 7, p <0.003), or *P. pelagicus* F5 (n = 3-6; t(6) = 7, p<0.004) at 625 ng.mL^-1^, indicating inhibition was not due to substrate limitations.

### 2.3 Heparin binding induces a conformational change in the Alzheimer’s disease beta secretase, BACE1

Hp binding has been proposed to occur at a location close to the active site of BACE1 [17], possibly within or adjacent to the prodomain sequence [48]. In-light of the contrasting and concentration dependant BACE1:GAG bioactivities, the ability of Hp and *P. pelagicus* F5 to induce structural changes in BACE1 has been investigated utilising circular dichroism (CD) spectroscopy at a range of w/w ratios; this also negates the intrinsic effect of the significant polydispersity for this class of biomolecules.

The CD spectra of BACE1 at pH 4.0 has previously been shown to contain a greater proportion of β-sheet and reduced α-helical content, compared to spectra obtained at pH 7.4, indicating that at an acidic pH, where BACE1 is most active, a conformational change can be observed by CD [52]. Consistent with this, the CD spectra of BACE1 in 50 mM sodium acetate buffer pH 4.0 (Figure 7 and 8), featured a positive peak at wavelengths below 200 nm, which can be attributed to a sum of α-helical and β- sheet structures [53]. The broad, negative band observed between wavelengths 250 - 200 nm, contains a peak at λ = 218 nm ∼ 208 nm, commonly associated with antiparallel β-sheets and α-helical structures, respectively [53] (Figure 7). The CD spectra of BACE1 at pH 4.0 can be estimated to have secondary structural composition of 9% α-helix, 31% antiparallel β-sheet, 16% turn and 44% other (NRMSD <0.1) when fitted against a library of representative proteins using BeStSel [50]. This was in close agreement with the BestSel secondary structure prediction based on x-ray crystallography of BACE1 at pH 4.0 (PDB accession no 2ZHS, [54] of of 7% α-helix, 30% antiparallel, 4% parallel, 12% turn and 47% other. Deviations between secondary structure predictions may be accounted for by subtle differences present within the BACE1 primary sequences.

**Figure 7:**
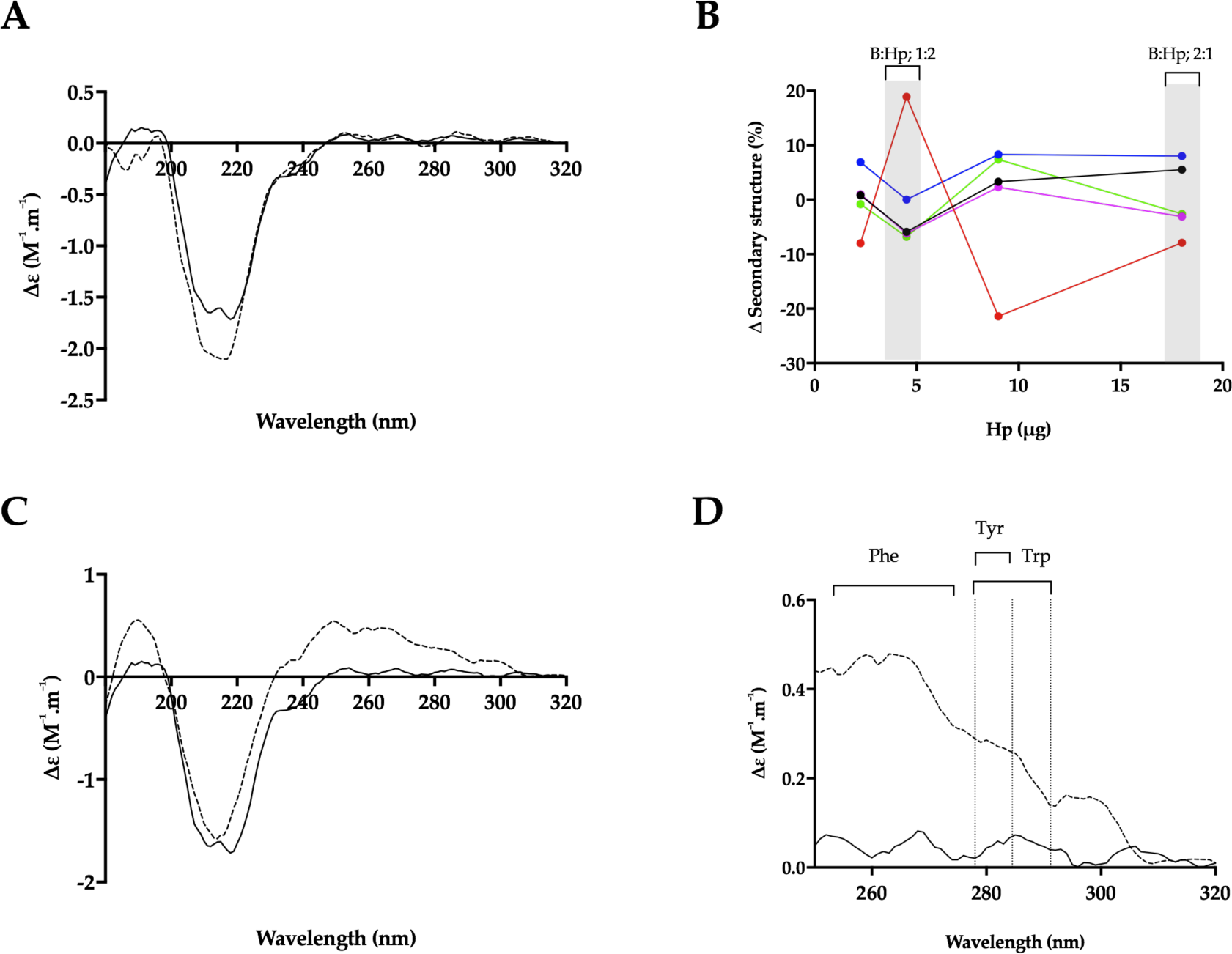
The structural change of BACE1 observed in the presence of Hp by circular dichroism (CD) spectroscopy. **(A)** CD spectra of BACE1 alone (solid line) or with Hp at a ratio of 1:2 (w/w; dashed line; B:Hp 1:2); **(B)** Δ secondary structure (%) of BACE1 upon the addition of increasing amounts of Hp; α-helix (black), antiparallel (red), parallel (blue), turn (magenta) and others (green) [50]. % structural change of B:Hp; 1:2 or 2:1 (w/w) ratio are highlighted in grey. **(C)** CD spectra of BACE1 alone (solid line) with Hp (dashed line) at a ratio of 2:1 w/w **(D)** Near-UV CD spectra of (C); respective absorption regions of aromatic amino acids are indicated [51]. Spectra were recorded in 50 mM sodium acetate buffer pH 4.0 in all panels.

**Figure 8:**
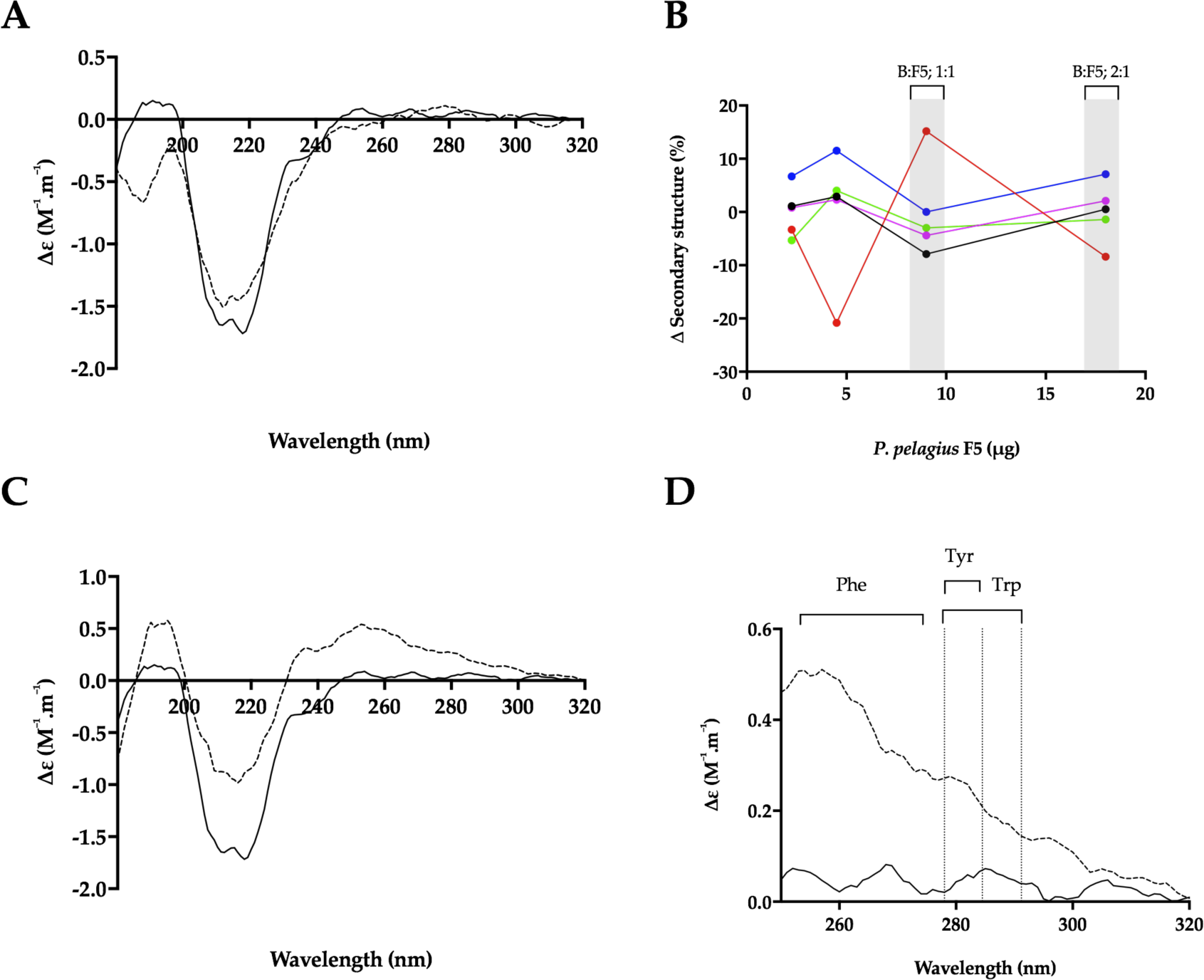
The structural change of BACE1 observed in the presence of *P. pelagicus* F5 by CD spectroscopy. **(A)** CD spectra of BACE1 alone (solid line) with *P. pelagicus* F5 (dashed line; ratio of 1:2 w/w; B:F5); **(B)** Δ secondary structure (%) of BACE1 upon the addition of increasing amounts of *P. pelagicus* F5; α-helix (black), antiparallel (red), parallel (blue), turn (magenta) and others (green) [50]. % structural change of B:F5; 1:2 or 1:1 ratio are highlighted in grey. **(C)** CD spectra of BACE1 alone (solid line) or with *P. pelagicus* F5 (dashed line; ratio of 1:1 w/w); **(D)** Near-UV CD spectra of (C); respective absorption regions of aromatic amino acids are indicated [51]. Spectra were recorded in 50 mM sodium acetate buffer pH 4.0 in all panels.

In the presence of a BACE1:Hp (B:Hp), ratio of 1:2 (w/w) where maximal inhibition was observed in FRET assays, the CD spectra of BACE1 exhibited increased negative ellipticity below λ = 222 nm, resulting in an estimated increase in α-helix (+ 6 %) and a reduction in antiparallel β-sheet (- 8 %) (NRMSD <0.1) [50] (Figure 7D). In comparison to Hp, BACE1 in the presence of *P. pelagicus* F5 (B:F5) at the same ratio (1:2; w/w), exhibited a slight increase in positive ellipticity between λ = 222 - 200 nm and decreases at λ < 200 nm, resulting in an estimated change in α-helical of + 1 % accompanied by a decrease in antiparallel β-sheet of 8 % (Figure 8D). This is in contrast to CD studies in the presence of peptide inhibitors, which did not reveal a secondary structural change in BACE1 [52].

The conformational change of BACE1 upon binding to Hp and *P. pelagicus* F5 was assessed over a range of ratios (Figure 7D and 8D). At a B:Hp ratio of 2:1 (w/w), a change in the characteristics of the CD spectrum of BACE1 was observed in the far-UV region (λ < 250 nm; Figure 7A) that was identified as a reduction in α-helix by 6% and an 19% increase in antiparallel β-sheet structures (NRMSD <0.1) [50]. In addition, an increase in positive ellipticity was observed in the near-UV region (250-300 nm; Figure 7B) following the addition of Hp, which may be attributed to a change in the tertiary structure of BACE1 involving aromatic amino acids [55,56] . In contrast, B:F5 at the same ratio of 2:1 (w/w), exhibited a decrease in ellipticity in the near- and far-UV region (λ < 300 nm; supplementary data).

The increase in positive ellipticity observed in the CD spectra of BACE1 in the near-UV region at B:Hp ratio of 2:1 (w/w), was also observed at a 1:1 (w/w) ratio of B:F5 (Figure 8B). The secondary structural change in the far-UV CD spectrum of BACE1 at a B:F5 ratio of 1:1 (w/w) between λ= 250 - 190 nm corresponded to a decrease in α-helix by 8% and an increase in antiparallel β-sheet structures by 15%.

### 2.4 Heparin and P. pelagicus F5 destabilise the Alzheimer’s BACE1

Both Hp and *P. pelagicus* F5 were shown to induce a conformational change in BACE1, in contrast to previous CD studies in the presence of peptide inhibitors [52]. Therefore, to explore whether the binding of Hp or *P. pelagicus* F5 alters the stability of BACE1, in a mechanism similar to known inhibitors, differential scanning fluorimetry (DSF) was employed to monitor the change in thermal stability. Previously identified BACE1 inhibitors have been shown to stabilize BACE1, exemplified by an increase in T_M_ values obtained through DSF measurements [57]. In the presence of a BACE1:Hp ratio of 1:2, a decrease in the T_M_ of BACE1 by 11°C was observed. In the presence of a BACE1:*P. pelagicus* F5 of 1:2 a decrease in the T_M_ of BACE1 was also observed by 10°C. The change in T_M_ of BACE1 induced by binding of either Hp or *P. pelagicus* F5 was not significantly different, (p = 0.1161 t= 2 df = 4). The destabilisation of BACE1 in the presence of both Hp and *P. pelagicus* F5 was found to be concentration dependent.

### 2.5 Attenuated anticoagulant activities of the *P. pelagicus* glycosaminoglycan extract

An important consideration when determining the therapeutic potential of a heparin-like polysaccharide against AD is the likely side effect of anticoagulation. Therefore, the prothrombin time (PT) and activated partial thromboplastin time (aPTT) were measured *P. pelagicus* F5 compared to Hp (193 IU.mg^-1^), in order to determine the overall effect on the extrinsic and intrinsic coagulation pathways, respectively (both assays also include the common coagulation pathway). In comparison to Hp, *P. pelagicus* F5 exhibited reduced anticoagulant activity in both the PT (Figure 9A; EC_50_ of 420.2 μg.mL^-1^ compared to 19.53 μg.mL^-1^, respectively) and aPTT (Figure 9B;EC_50_ 43.21 μg.mL^-1^ compared to 1.66 μg.mL^-1^, respectively) coagulation assays. Both results show that the extract presents a negligible anticoagulant activity.

**Figure 9:**
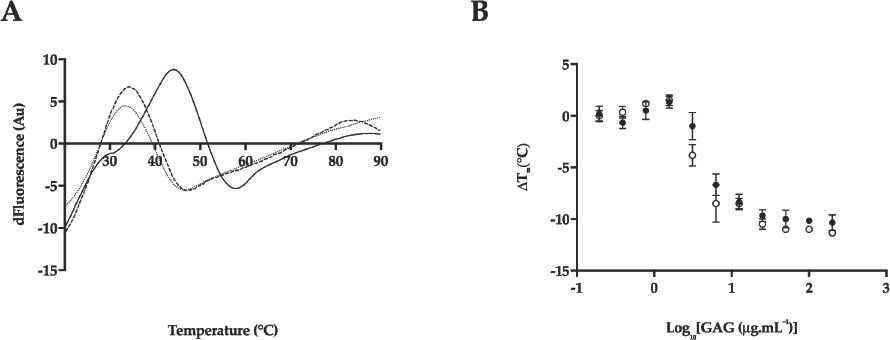
**(A)** First differential of the DSF thermal stability profile of BACE1 alone (1 μg; solid line), and with Hp (2 μg; dotted line) or *P. pelagicus* F5 (2 μg; dashed line), in 50 mM sodium acetate, pH 4.0; **(B)** Δ T_m_ of BACE1 with increasing [Hp] or [*P. pelagicus* F5] (open or closed circles, respectively).

**Figure 10:**
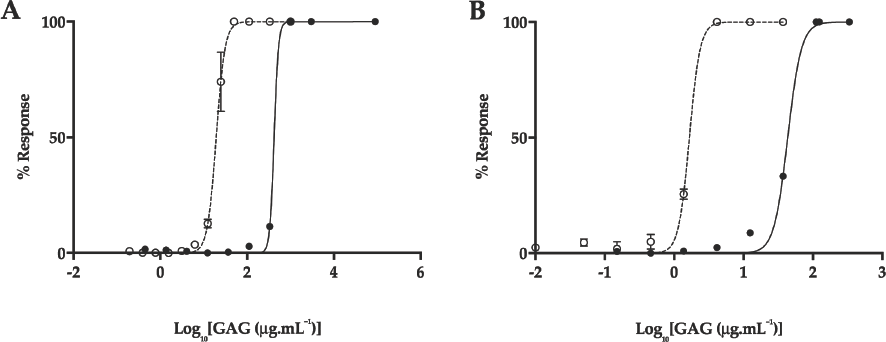
**(A)** Prothrombin time (PT) and **(B)** activated partial thromboplastin time (aPTT) inhibitory response 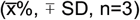 for Hp (open circle, dashed line) and *P. pelagicus* F5 (closed circle, solid line);. ***PT:*** Hp EC_50_ = 19.53 μg.mL^-1^; *P. pelagicus* F5, EC_50_ = 420.2 μg.mL^-1^; ***aPTT***: Hp EC_50_ = 1.66 μg.mL^-1^;*P. pelagicus* F5, EC_50_ = 43.21 μg.mL^-1^.

## 3. Discussion

The glycosaminoglycan extract isolated from *P. pelagicus* was observed to possess similar electrophoretic behavior to mammalian HS and Hp, with no bands identified corresponding to CS or DS standards. In contrast, the FTIR and HSQC analyses of *P. pelagicus* F5 identified regions corresponding to both HS and CS saccharides within the extract. PCA analysis of the FTIR spectra revealed *P. pelagicus* F5 contained features associated with both HS/Hp and CS/DS, which are typical of crude heparin preparations [44]. This was confirmed by HSQC NMR which identified N-acetyl peaks associated with both galactosamine (CS) and glucosamine (HS). The absence of an IdoA signal from the NMR spectra suggests *P. pelagicus* F5 resembles HS and CS more closely than DS/Hep [58,59]. Peaks corresponding to Gal-6S and 6-OH were identified by NMR analysis, with no detectable 4-O-sulfation, indicating that the CS component of *P. pelagicus* F5 resembles CSC saccharides. The HS component possesses > 70% N-sulfated moieties, which is greater than mammalian HSs published previously, but is not as heavily N-sulfated as mammalian heparins. An intermediate proportion of trisulfated Δ-disaccharides were also identified post bacterial lyase digestion in *P. pelagicus* F5, when compared to mammalian HS and Hp samples. Furthermore, the *P. pelagicus* F5 extract contained a low proportion of Δ-UA-GlcNAc/ Δ-UA-GlcNS, which is typical of more heparin-like structures. This suggests that the HS/Hp component of *P. pelagicus* F5 consists of a hybrid structure lacking the domain structure of HS and the highly sulfated regions of Hp.

The absence of a band migrating in a similar manner to that of CS when *P. pelagicus* F5 was subjected to agarose gel electrophoresis suggests that the polysaccharide is not a mixture of HS and CS chains. The simplicity of the signals in the HSQC spectrum suggests either two separate populations, or two distinct domains, while the former is not consistent with the agarose gel electropherogram mentioned previously. The PCA of the FTIR spectra is also in agreement with the presence of discrete, rather than mixed HS/CS sequences. The precise nature of the arrangement of these stretches remains unknown, although it is well documented that marine-derived GAGs harbour significant and unusual structural features, when compared to those present within their mammalian counterparts [25,26,38,60–69]. Studies to resolve this technically demanding question are currently in progress.

The *P. pelagicus* F5 extract was found to possess significant inhibitory potential against human BACE1, in a manner akin to that of mammalian Hp, as demonstrated by comparable IC_50_ concentrations determined via FRET. The ability of *P. pelagicus* F5 to promote BACE1 bioactivity at lower concentrations, owing to the presence of the BACE1 pro-domain [48,49] appears to be at a diminished level compared to mammalian Hp, suggesting differences between these GAGs and the nature of their interactions with human BACE1. This was exemplified when the secondary structural changes in BACE1 (evident from CD) in the presence of Hp or *P. pelagicus* F5 were examined.

BACE1 has previously been observed to adopt a unique secondary structure at pH 4.0, where catalytic activity is increased, resulting in a predicted increase in beta-sheet and a reduction in alpha-helical structures [52]. When the changes in the secondary structure (evident from CD) of BACE1 in the presence of high concentrations of Hp (BACE1:Hp ratio of 1:2) was examined a shift towards the structural features observed for BACE1 alone at pH 7.4. was observed (increase in alpha-helical and reduction in beta sheets). At high concentrations (B:F5 ratio of 1:2), the *P. pelagicus* F5 extract induced similar, but not identical, changes to the secondary structure of human BACE1, when compared to those of Hp at the same ratio.

In contrast, the CD spectra observed for B:Hp complexes under conditions that facilitate BACE1 promotion (i.e. low Hp concentrations), demonstrated evidence of an interaction that involves the aromatic amino acids (near UV CD). Tyr-71 is located within the BACE1 flap that has previously been identified to change conformation between the flap-open and flap-closed states [70] . Unfortunately, due to the location of the aromatic residues on the surface of the protein, it is not possible to conclude definitely whether interaction(s) of Hp-based inhibitors with human BACE1 occur at, or near to, the active site. This interpretation is consistent with the previous reports that a conformational change in BACE1 may occur upon heparin binding, which would be required to allow access into the active site [48] . In addition, the increase in BACE1 activity by heparin has been shown to be followed by BACE1 inhibition [49], which may suggest this arrangement is required to allow access to the active site. The results also support the work by Scholefield *et al.* [17] who showed that the mode of Hp inhibition is non-competitive, and can prevent access of the substrate.

At lower GAG concentrations, differences in BACE1 secondary structure were observed between the B:Hp and B:F5 complexes in the CD spectra, although a similar change in the near UV CD spectra of BACE1 was observed with increased amounts of *P. pelagicus* F5. This may be accounted for by the reduced potency of *P. pelagicus* F5 with regard to activating BACE1, or indicative of an alternative interaction. The conformational change induced in the near-UV CD spectra of BACE1 is solely the result of the HS/Hp-like component of the *P. pelagicus* F5 extract. CS has previously been shown to possess diminished BACE1 promotion activity compared to Hp/HS [48].

From a mechanistic standpoint, the decrease in the T_m_s observed using DSF for both the human BACE1 protein in the presence of either Hp or the *P. pelagicus* F5 extract, when compared to human BACE1 alone, suggests that the mode of BACE1 inhibition by this class of carbohydrates could both involve structural destabilisation. The Hp-induced thermal instability of human BACE1 occurs in a concentration dependent manner, akin to that of the inhibitory potential of Hp in the FRET-based bioactivity assay. As for the FRET-based, BACE1 inhibition assays, *P. pelagicus* (F5) also induces comparable destabilisation of BACE1 with similar T_m_ values. A graph of BACE1:GAG T_m_ vs concentration demonstrates similar profiles for the *P. pelagicus* GAG extract and that of mammalian Hp. The relationship between Hp and *P. pelagicus* F5 concentration and biological properties that coexists for both FRET-based, BACE1 inhibition and DSF is not mirrored at defined concentrations of Hp and *P. pelagicus* F5 with regard to their distinct CD spectra and predicted secondary structure. This would suggest complex and distinct modes of interactions are present.

One of the major obstacles that precludes the use of mammalian Hp compounds as potential BACE1 inhibitors and pharmaceutical candidates in general, is that of the significant anticoagulant potential residual within the biomolecule. This anticoagulant potential is afforded by the propensity of Hp to interact with antithrombin and thereby inhibit the human coagulation pathway, which unperturbed, ultimately results in fibrin clot formation. The anticoagulant potential of *P. pelagicus* F5 has been shown to be highly attenuated in contrast to mammalian Hp, as measured by both the aPTT and PT clotting assays. These coagulation assays are routinely employed, in clinical settings, to screen for the common pathway in combination with either the intrinsic (apTT) or extrinsic pathways (PT).

Major limitations for the repurposing of Hp from mammalian origins include the potential contamination risk from animal-derived viruses or prions, notably bovine spongiform encephalopathy as well as differing cultural and religious mores throughout the world. Sourcing a GAG-based inhibitor of BACE1 from marine origins would lessen the risks associated with use of mammalian Hp as it is not animal-derived and will be free from contamination with mammalian pathogens. In addition, the abundance of processed marine waste available as a byproduct of the food industry [26. 64] offers a novel and valuable resource for the large-scale isolation of GAG-like polysaccharides. These marine-sourced polysaccharides have a significant potential for future therapeutic applications [71–73].

## 4. Materials and Methods

### 4.1 Extraction of glycosaminoglycans from *Portunus pelagicus*

2.4 kg of *Portunus pelagicus* tissue (Yeuh Chyang Canned Food Co., Ltd., Vietnam) was homogenised with excess acetone (VWR, UK) and agitated for 24 hours at r.t. Defatted, *P. pelagicus* tissue was recovered via centrifugation, 5,670 rcf at r.t. for 10 minutes and allowed to air dry. The tissue was then subjected to extensive proteolytic digestion (Alcalase®; Novozymes, Denmark) using 16.8 U.kg^-1^ of dried tissue mass, in PBS (w/v; Gibco, UK) made up to a final concentration of 1 M NaCl (Fisher Scientific, UK), pH 8.0 and incubated at 60°C for 24 hours. Post digestion, the supernatant was collected via centrifugation (5,670 g for 10 minutes, r.t.), and subjected to ion exchange chromatography employing Amberlite IRA-900 resin (Sigma-Aldrich, UK; hydroxide counterion form) for 24 hours under constant agitation at r.t. Ion exchange resin was recovered by filtration and washed successively with distilled H_2_O (Fisher Scientific, UK) at 60°C with two volumes of water and 10 volumes of 1 M NaCl at r.t. The ion exchange resin was then re-suspended in 1 L of 3 M NaCl and agitated for 24 hours at r.t. The ion exchange resin was removed and the filtrate added to ice cold methanol (VWR, UK), 1:1 (v/v) prior to incubation for 48 hours at 4°C. The precipitate formed was recovered by centrifugation at 4°C, 15,400 g for 1 hour and re-suspended in distilled H_2_O. The crude *P. pelagicus* extract was dialysed against distilled H_2_O (3.5 kDa MWCO membrane; Biodesign, USA) for 48 hours prior to syringe filtration (0.2 µm) and lyophilisation. The crude GAG extract was re-suspended in 1 mL of HPLC grade water and loaded onto a pre-packed DEAE-Sephacel column (10 mm I.D. x 10 cm; GE Healthcare, UK) at a flow rate of 1 mL.min^-1^. The column was eluted using a stepwise NaCl gradient of 0, 0.25, 0.5, 0.8, 1 and 2 M NaCl at a flow rate of 1 mL.min^-1^, with elution monitored in-line at λ_abs_ = 232 nm (using a UV/Vis, binary gradient HPLC system; Cecil Instruments, UK), resulting in six fractions (F1 - F6, respectively). Each of the eluted fractions were dialysed against distilled H_2_O, employing a 3.5 kDa MWCO (Biodesign, USA) for 48 hours under constant agitation. The retentate obtained for F5 was lyophilised and stored at 4°C prior to use.

### 4.2 Agarose gel electrophoresis

Agarose gel electrophoresis was performed in 0.55% (w/v) agarose gels (8 × 8 cm, 1.5 mm thick) prepared in 1,3-diaminopropane-acetate buffer pH 9.0 (VWR, UK), 2-7.5 μg of the of *P. pelagicus* F5 or GAG standards were subjected to electrophoresis utilizing a X-Cell SureLock™ Mini-Cell Electrophoresis System (ThermoFisher, UK). Electrophoresis was performed in 0.5 M 1,3-diaminopropane-acetate buffer (pH 9.0), at a constant voltage of 150 V (∼100 mA) for ∼30 minutes or until the dye front had migrated ∼ 8 cm from the origin. The gels were then precipitated with 0.1% w/v cetyltrimethylammonium bromide solution (VWR, UK) for a minimum of 4 hours and then stained for 1 hour in 0.1% toluidine blue dissolved in acetic acid:ethanol:water (0.1:5:5). Gels were de-stained in acetic acid:ethanol:water (0.1:5:5 v/v) for ∼ 30 minutes prior to image acquisition with GIMP software and processing with ImageJ.

### 4.3 Attenuated FTIR spectral analysis of marine-derived glycosaminoglycans

Samples (10 mg, lyophilised) were recorded using a Bruker Alpha I spectrometer in the region of 4000 to 400 cm^-1^, for 32 scans at a resolution of 2 cm^-1^ (approx 70 seconds acquisition time), 5 times. Spectral acquisition was carried-out using OPUS software (Bruker) with correction to remove the residual spectrum of the sampling environment.

Spectral processing and subsequent data analyses were performed using an Asus Vivobook Pro (M580VD-EB76), equipped with an intel core i7-7700HQ. Spectra were smoothed, employing a Savitzky-Golay smoothing algorithm (R studio v1.1.463; *signal* package, *sgolayfilter)*, to a 2^nd^ degree polynomial with 21 neighbours prior to baseline correction employing a 7th order polynomial and subsequent normalisation (0-1). CO_2_ and H_2_O regions were removed prior to further analysis, in order to negate environmental variability (< 700 cm^-1^, between 2000 and 2500 cm^-1^ and >3600 cm^-1^). Second derivatives plots were calculated using the Savitzky-Golay algorithm, with 41 neighbours and a 2^nd^ order polynomial.

The normalised and corrected matrix of intensities was subject to PCA using singular value decomposition in R studio (v1.1.463) with the mean centred, *base prcomp* function deployed.

### 4.4 Nuclear Magnetic Resonance (NMR)

NMR experiments were performed upon *P. pelagicus* F5 (5 mg) dissolved in D_2_O (600 µL; VWR, Brazil) containing TMSP (0.003% v/v; VWR, Brazil) at 343 K using a 500 MHz Avance Neo spectrometer fitted with a 5 mm TXI Probe (Bruker). In addition to 1-dimensional (^1^H) spectra, ^1^H-^13^C Heteronuclear Single-Quantum Correlation (HSQC) 2-dimensional spectra were collected using standard pulse sequences available. Spectra were processed and integrated using TopSpin (Bruker).

### 4.5 Constituent Δ-disaccharide analysis of Hp/HS-like, marine-derived carbohydrates

Hp and *P. pelagicus* F5 carbohydrate samples were re-suspended in lyase digestion buffer (50 µL; 25 mM sodium acetate, 5 mM calcium acetate (VWR, UK), pH 7) prior to exhaustive digestion by the sequential addition of a cocktail of the three recombinantly expressed heparinase enzymes (I, III & II) from the soil bacterium *Flavobacterium heparinum* (2.5 mIU.mg^-1^; Iduron, UK). Samples were incubated for 4 hrs at 37 °C prior to a further addition of the same quantity of enzymes and an additional overnight incubation. Samples were then heated briefly at 95°C post enzyme digestion (5 mins) and allowed to cool.

Denatured heparinase enzymes were removed from the sample solution by immobilisation upon a pre-washed (50% methanol (aq.) followed by HPLC grade H_2_O) C^18^ spin column (Pierce, UK); whereby the newly liberated Δ-disaccharides were present in the column eluate upon washing with HPLC grade H_2_O.

Lyophilised Δ-disaccharide samples from Hp and *P. pelagicus* F5 were desalinated by immobilisation up on graphite spin columns (Pierce, UK) that had been extensively prewashed with 80% acetonitrile, 0.5% (aq.) trifluoroacetic acid and HPLC grade H_2_O prior to use. Δ-disaccharides liberated from the exhaustive, heparinase digestion were separated from buffer salts by extensive washing with HPLC grade H_2_O prior to elution with a solution of 40% acetonitrile, 0.5% trifluoroacetic acid (aq.). Contaminant, non Δ-disaccharide components of the spin column eluate were removed by serial lyophilization prior to chromatographic separation, using high performance anion exchange chromatography (HPAEC).

Heparinase digested samples (50 µg) were made up in HPLC grade H_2_O (1 mL) immediately before injection onto a ProPac PA-1 analytical column (4 × 250 mm, ThermoFisher Scientific, UK), pre-equilibrated in HPLC grade H_2_O at a flow rate of 1 mL.min^-1^. The column was held under isocratic flow for 10 mins prior to developing a linear gradient from 0 to 2 M NaCl (HPLC grade; VWR, UK) over 60 mins. Eluted Δ-disaccharides were detected absorbing within the UV range (λ_abs_ = 232 nm) via the unsaturated C=C bond, present between C_4_ and C_5_ of the uronic acid residues, introduced as a consequence of lyase digestion.

Authentic Δ-disaccharide reference standards, comprising the 8 common standards found in Hp and HS (Iduron, UK), were employed as a mixture (each at 5 μg.mL^-1^) and served as a chromatographic references with elution times cross-correlated with Hp and *P. pelagicus* F5 samples. The column was washed extensively, with 2 M NaCl and HPLC grade water, prior to use and between runs.

### 4.6 Determination of human BACE1 inhibitory activity using Förster resonance energy transfer

*P. pelagicus* F5 and Hp were assayed for inhibitory potential against human beta-secretase (BACE1) using the fluorescence resonance energy transfer (FRET) inhibition assay, essentially as described by Patey et al (2006) [18]. Human BACE1 (312.5 ng), and *P. pelagicus* F5 or Hp were incubated in 50 mM sodium acetate pH 4.0 at 37°C for 10 minutes, followed by the addition a quenched fluorogenic peptide substrate (6.25 μM; Biomatik, Canada; MCA-SEVNLDAEFRK(DNP)RR-NH_2_; pre-incubated at 37°C for 10 minutes) to a final well volume of 50 μL. Fluorescent emission was recorded using a Tecan Infinite® M200 Pro multiwell plate reader with i-control™ software (λ_ex_ = 320 nm, λ_em_ = 405 nm) over 90 minutes. The relative change in fluorescence per minute was calculated in the linear range of the no inhibitor control, with normalized percentage inhibition calculated (% ± SD, n = 3) compared to the 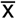 of substrate only and no inhibitor control, followed by fitting to a four-parameter logistics model using Prism 7 (GraphPad).

### 4.7 Secondary structure determination of human BACE1 by circular dichroism spectroscopy

The circular dichroism (CD) spectra of native, human BACE1 (6.12 μM, 30 μl; Acro Biosystems, USA) in 50 mM sodium acetate (pH 4.0; VWR, UK) was obtained using a J-1500 Jasco CD spectrometer and Spectral Manager II software, equipped with a 0.2 mm path length quartz cuvette (Hellma, USA) operating at a scan speed of 100 nm.min^-1^ with 1 nm resolution over the range λ = 190 - 320 nm. Spectra obtained were the mean of five independent scans. Human BACE1 was buffer exchanged (in order to remove commercially supplied buffer) prior to spectral analysis using a 10 kDa Vivaspin centrifugal filter (Sartorius, Germany) at 12,000 g washed thrice. Collected data was processed using Spectral Manager II software and data analysis carried out with GraphPad Prism 7, employing a second order polynomial smoothed to 9 neighbours. Secondary structure prediction was performed utilizing the BeStSel analysis server on the unsmoothed data [50].To ensure the CD spectral change of BACE1 in the presence of each GAG was not altered owing to the addition of the GAG alone, which are known to possess CD spectra at high concentrations [74,75], GAG control spectra were subtracted before analysis. In addition, the theoretical, summative CD spectra was confirmed to differ from the observed experimental CD spectra, thereby indicating that the change in the CD spectra compared to that of BACE1 alone is a result of a conformational change upon binding to the GAG. The conformational change observed is believed to occur as a result of changes solely in BACE1 secondary structure, as GAG controls exhibited negligible spectra at the concentration used. All CD data have been presented with GAG controls subtracted.

### 4.8 Investigating the thermal stability of human BACE1 with differential scanning fluorimetry

Differential scanning fluorimetry (DSF) was carried out using the method of Uniewicz et al. (2014) [76] based on a modification to the original method of Niesen et al. (2007) (Uniewicz, Ori, Ahmed, Yates, & Fernig, 2014) [76,77]. DSF was performed on human BACE1 (1 μg) using 96-well qPCR plates (AB Biosystems, UK) with 20x Sypro Orange (Invitrogen, UK) in 50 mM sodium acetate, pH 4.0 in a final well volume of 40 μl. Hp or mGAG were included, as necessary, to a maximal concentration of 200 μg.mL^-1^. An AB Biosystems StepOne plus qPCR machine, with the TAMRA filter set deployed, was used to carry out melt curve experiments, with an initial incubation phase of 2 minutes at 20°C, increasing by 0.5°C increments every 30 seconds up to a final temperature of 90°C. Data analysis was completed using Prism 7 (GraphPad) with first derivative plots smoothed to 19 neighbours, using a second order polynomial (Savitzky-Golay). The peak of the first derivatives (yielding T_m_s) was determined using MatLab (MathWorks) software.

### 4.9 Activated partial thromboplastin time (APTT)

Serially diluted GAG samples (25 μl) were incubated with pooled, normal human citrated plasma (50 μl; Technoclone, UK) and Pathromtin SL reagent (50 μl; Siemens, UK) for 2 mins at 37°C prior to the addition of calcium chloride (25 μl, 50 mM; Alfa Aesar, UK). The time taken for clot formations to occur (an upper maximal of 2 mins was imposed, represented as 100% inhibition of clotting) were recorded using a Thrombotrak Solo coagulometer (Axis-Shield). HPLC grade H_2_O (0% inhibition of clotting, representing a normal aPTT clotting time, of ≈ 37-40 seconds) and porcine mucosal heparin (193 IU.mg^-1^ ; Celsus, USA) were screened as controls. The EC_50_ values of all test and control samples were determined using a sigmoidal dose response curve fitted with GraphPad Prism 7.

### 4.10 Prothrombin time (PT)

Serially diluted GAGs (50 μl) or control (H_2_O, HPLC grade) were incubated with pooled, normal human citrated plasma (50 μl) for 1 minute at 37°C prior to the addition of Thromborel S reagent (50 μl; Siemens, UK). The time taken for clot formations to occur (an upper maximal of 2 minutes was imposed, representing 100% inhibition of clotting) were recorded using a Thrombotrak Solo coagulometer. HPLC grade H_2_O (0% inhibition of clotting, representing a normal PT clotting time of ≈ 13-14 seconds) and porcine mucosal heparin (193 IU.mg^-1^; Celsus, USA) were screened as controls. The EC_50_ values of all test and control samples were determined using a sigmoidal dose response curve fitted with GraphPad Prism 7.

## Supplementary Materials

Figure S1: The CD structural change of BACE1 observed in the presence of *P. pelagicus* F5 with a ratio of 2:1 w/w.

## Acknowledgments

The authors would like to thank Dr. Sarah Taylor for technical assistance with the use of CD instrumentation.

## Supplementary Materials

**Supplementary 1:**
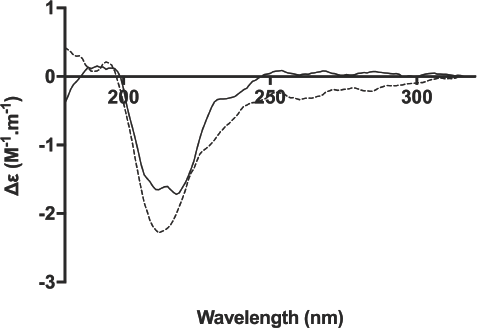
The CD structural change of BACE1 (solid) observed in the presence of *P. pelagicus* F5 (dashed) with a ratio of 2:1 w/w, B:F5.

